# Reconstructing feedback representations in ventral visual pathway with a generative adversarial autoencoder

**DOI:** 10.1101/2020.07.23.218859

**Authors:** Haider Al-Tahan, Yalda Mohsenzadeh

## Abstract

While vision evokes a dense network of feedforward and feedback neural processes in the brain, visual processes are primarily modeled with feedforward hierarchical neural networks, leaving the computational role of feedback processes poorly understood. Here, we developed a generative autoencoder neural network model and adversarially trained it on a categorically diverse data set of images. We hypothesized that the feedback processes in the ventral visual pathway can be represented by reconstruction of the visual information performed by the generative model. We compared representational similarity of the activity patterns in the proposed model with temporal (magnetoencephalography) and spatial (functional magnetic resonance imaging) visual brain responses. The proposed generative model identified two segregated neural dynamics in the visual brain. A temporal hierarchy of processes transforming low level visual information into high level semantics in the feedforward sweep, and a temporally later dynamics of inverse processes reconstructing low level visual information from a high level latent representation in the feedback sweep. Our results append to previous studies on neural feedback processes by presenting a new insight into the algorithmic function and the information carried by the feedback processes in the ventral visual pathway.

**Author summary:** It has been shown that the ventral visual cortex consists of a dense network of regions with feedforward and feedback connections. The feedforward path processes visual inputs along a hierarchy of cortical areas that starts in early visual cortex (an area tuned to low level features e.g. edges/corners) and ends in inferior temporal cortex (an area that responds to higher level categorical contents e.g. faces/objects). Alternatively, the feedback connections modulate neuronal responses in this hierarchy by broadcasting information from higher to lower areas. In recent years, deep neural network models which are trained on object recognition tasks achieved human-level performance and showed similar activation patterns to the visual brain. In this work, we developed a generative neural network model that consists of encoding and decoding sub-networks. By comparing this computational model with the human brain temporal (magnetoencephalography) and spatial (functional magnetic resonance imaging) response patterns, we found that the encoder processes resemble the brain feedforward processing dynamics and the decoder shares similarity with the brain feedback processing dynamics. These results provide an algorithmic insight into the spatiotemporal dynamics of feedforward and feedback processes in biological vision.

## Introduction

In just a couple of hundred milliseconds, our brain interprets the visual scene around us [1–5], identifies faces [6, 7], recognizes objects [8–13], and localizes them [14–18]. Decades of cognitive neuroscience research has demonstrated that the brain accomplishes these complicated tasks through a cascade of hierarchical processes in the ventral visual stream starting in the early visual cortex (EVC) and culminating in the inferior temporal (IT) cortex.

While the feedforward recruitment of this hierarchy explains the core neural response patterns underlying visual recognition [19–21], it is unable to account for behavioural and neural dynamics observed in years of psychophysical, neurophysiological, and magneto/electrophysiological experiments [12, 22–25]. Indeed, variable timing of neural responses to visual stimuli beyond 200ms has been frequently associated with accumulation of sensory evidence through recurrent processes in the visual brain. However, the precise computational role of neural recurrent/feedback processes remains poorly understood at the system level. In particular, the algorithmic function of feedback processes and the type of information sent back along the visual hierarchy is still unknown.

To address this question, we develop a generative model which is adversarially trained on a diverse set of image categories. The model consists of two sub-networks: (i) An encoder sub-network receives a given visual stimulus, processes it in a hierarchy of neural layers to eventually produce a latent representation (code) of the visual input and (ii) a decoder sub-network which receives the latent representation and aims to reproduce the visual input from the information encoded in the latent representation.

This generative model enables us to not only investigate the encoding process of visual representations along the hierarchy of the encoder sub-network layers but also provides us an insight into the reverse process i.e. reconstructing the representations along the decoder sub-network layers. We hypothesize that the visual information in the encoder sub-network in our computational model mimics the feedforward pathway in the ventral visual stream and the decoder sub-network which performs the reverse function may reveal the representations along the feedback pathway. To test this hypothesis, after training the proposed model, we compare the representations along its layers with magnetoencephalography (MEG) and functional magnetic resonance imaging (fMRI) data acquired from fifteen human participants in a visual recognition experiment [13].

Our model identified two separate dynamics of representational similarities with MEG temporal data. The first one is consistent with the temporal hierarchy of processes transforming low level visual information into high level semantics in the feedforward sweep, and the second one reveals a temporally subsequent dynamics of inverse processes reconstructing low level visual information from a high level latent representation in the feedback sweep.

Further, comparison of encoder and decoder representations with two fMRI regions of interests, namely EVC and IT, revealed a growing categorical representation along the encoder layer (feedforward sweep) similar to IT and a progression in detail visual representations along the decoder layers (feedback sweep) akin to EVC.

## Results

### Construction of a generative model performing image reconstruction

Previous work revealed that deep convolutional neural networks (DNNs) trained on image classification develop hierarchical representations similar to the cascade of processes along ventral visual pathway [3, 26–33]. However, neuroscience evidence suggests top-down modulations of neural responses which occur after some delay through abundant number of feedback connections in visual cortex are critical to resolving visual recognition in the brain [12, 23, 24]. Therefore, these feedforward deep neural network models do not fully represent the complex visual processes in the ventral visual pathway. Here, we investigate whether a deep generative model trained to compress and reconstruct images could reveal similar representations as feedforward and feedback processes in the ventral visual pathway. With this aim, we developed a deep generative autoencoder neural network model using adverserial autoencoder (AAE) framework [34]. AAE is a generative adverserial network (GAN) [35] where the generator has an autoencoder architecture. Figure 1 depicts our proposed model architecture. The autoencoder generator consists of two main components: 1) an encoder which receives the visual stimuli and performs a cascade of simple operations such as convolution, pooling, and normalization to map the visual input to a latent feature vector (LV); 2) a decoder which receives the latent vector and performs a cascade of simple deconvolution, upsampling and normalization operations to reconstruct the input visual stimuli from information encoded in the latent space. The model is trained with two objectives - a reconstruction loss criterion, and an adversarial criterion. The dual objectives training turns the autoencoder into a generative model whose latent space learns data distribution properties that enables generative process and avoids overfitting to the reconstruction objective. We hypothesize that the encoder sub-network models the feedforward pathway of processes in ventral visual stream, while the decoder sub-network models the reconstruction of visual features in the feedback pathway.

**Fig 1.**
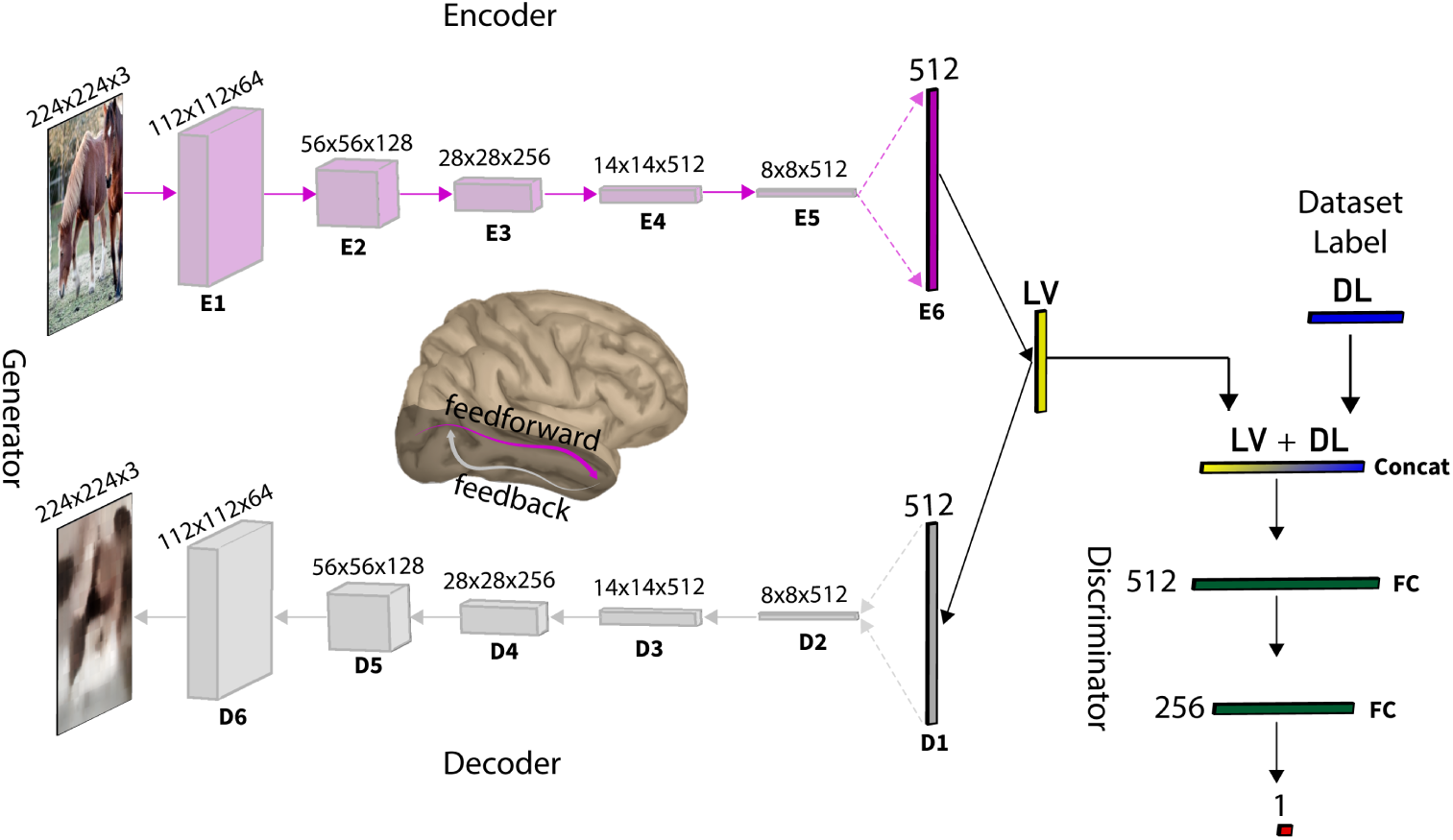
Computational model architecture. The model is a generative adverserial network. The generator is an autoencoder consisting of five convolutional blocks (E1-E5) and one fully connected layer (E6) in the encoder and one fully connected layer (D1) followed by five deconvolutional blocks in the decoder (D2-D6). Each convolutional block encompasses batch normalization, convolution, nonlinear activation function, and pooling operations. Alternatively, each deconvolutional block encompasses batch normalization, deconvolution, nonlinear activation function, and upsampling operations. The discriminator consists of two fully connected layers. The training Data set consists of 1,980,000 images organized into four super-ordinate categories: (i) Faces, (ii) Animates, (iii) Objects, (iv) Scenes. **LV** denotes the latent vector generated by the encoder and **DL** is a one-hot data set label (one of the four mentioned training data sets). Both vectors are concatenated and fed to the discriminator, while only the latent vector is fed to the decoder.

To train our model, we assembled a super category data set (see Materials and methods section for details). The super category data set includes 1,980,000 images from four equally distributed categories of (1) Faces, (2) Animates, (3) Objects, and (4) Scenes. The rational behind assembling and using this data set is two-fold: (1) ecologically, the human brain learns to develop high-level category representations across multiple recognition tasks (e.g. faces, animals, objects, scenes, etc.); Indeed, years of neuroscience research have identified a cascade of brain regions along ventral visual stream starting in early visual cortex (EVC) which processes low level visual features and eventually resolves visual categories at the end of the hierarchy in inferior temporal cortex (IT). (2) In this study, we compare model representations with brain imaging data (fMRI and MEG) from a visual recognition experiment [13]. We maintained consistency with the four categories from the stimulus set utilized in the brain imaging experiment which includes 156 images. Please note that this stimulus set was not used for training the model. After training the model on super category data set for over 800 epochs when the adverserial and reconstruction losses reached their local minima on the training set (Figure 2A), we determined how well our model performed on the 156 image set (as a testing dataset). Figure 2B illustrates this image set and the corresponding reconstructed images. The model performed the reconstruction on the training dataset with 0.109 ± 0.006 mean absolute error (MAE) and on the 156 image set with 0.119 ± 0.003 MAE between the input image and the corresponding reconstructed image. We computed the upper-bound performance of the model by sampling random pairs of images from the training and testing sets performing MAE between the different image pairs. On the training set, we observed a MAE of 0.53 ± 0.014 between the random image pairs and 0.62 ± 0.017 on the random pairs from the testing set. These results show that the model not only have converged to an optimum but also generalizes well to the testing set.

**Fig 2.**
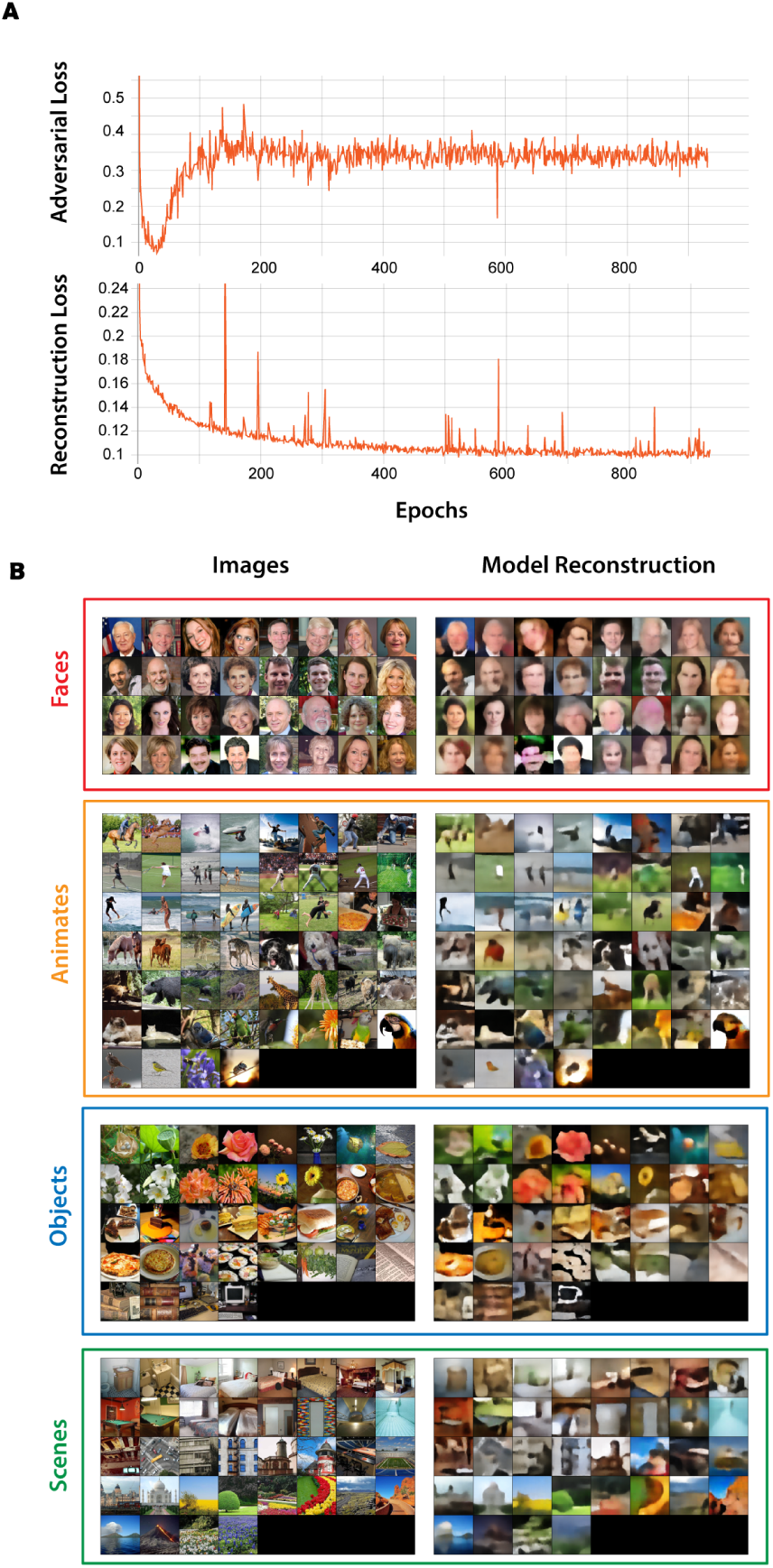
Computational model performance. (A) Adversarial and Reconstruction loss over epochs. (B) 156 image set and their corresponding reconstructed image by the model.

### Representational similarity of the generative model to early and late brain regions in the ventral visual stream

We first determined the encoder/decoder representational similarities with early and late brain regions along the ventral visual cortex. For this, we chose early visual cortex (EVC) and inferior temporal cortex (IT) defined anatomically based on [36] and employed the representational similarity analysis (RSA) method [37, 38] as the integrative framework for model-brain comparisons.

For each region of interest (ROI), we extracted the fMRI response patterns to each image, vectorized it and computed condition-specific pairwise distances (1-Pearson’s R) to create a 156 × 156 representational dissimilarity matrix (RDM) per participant. We also fed the images to the generator and extracted layer-specific activations for each image condition. Then by computing the pairwise distances (1-Pearson’s R) of image evoked layer activation patterns, we created layer-specific RDMs (see Figure 3 and Materials and methods section for details). We then compared subject-specific ROI RDMs with the model layer RDMs by computing Spearman’s correlations (Figure 3ADE). Figure 4A and B show subject averaged RDMS and their corresponding 2-dimensional multidimensional scaling (MDS) visualizations of EVC and IT, respectively. As expected EVC shows a random pattern across categories, whereas IT demonstrates clear categorical distinctions. Figure 4C-D compares the encoder and decoder layers correlations with EVC and IT RDMs.

**Fig 3.**
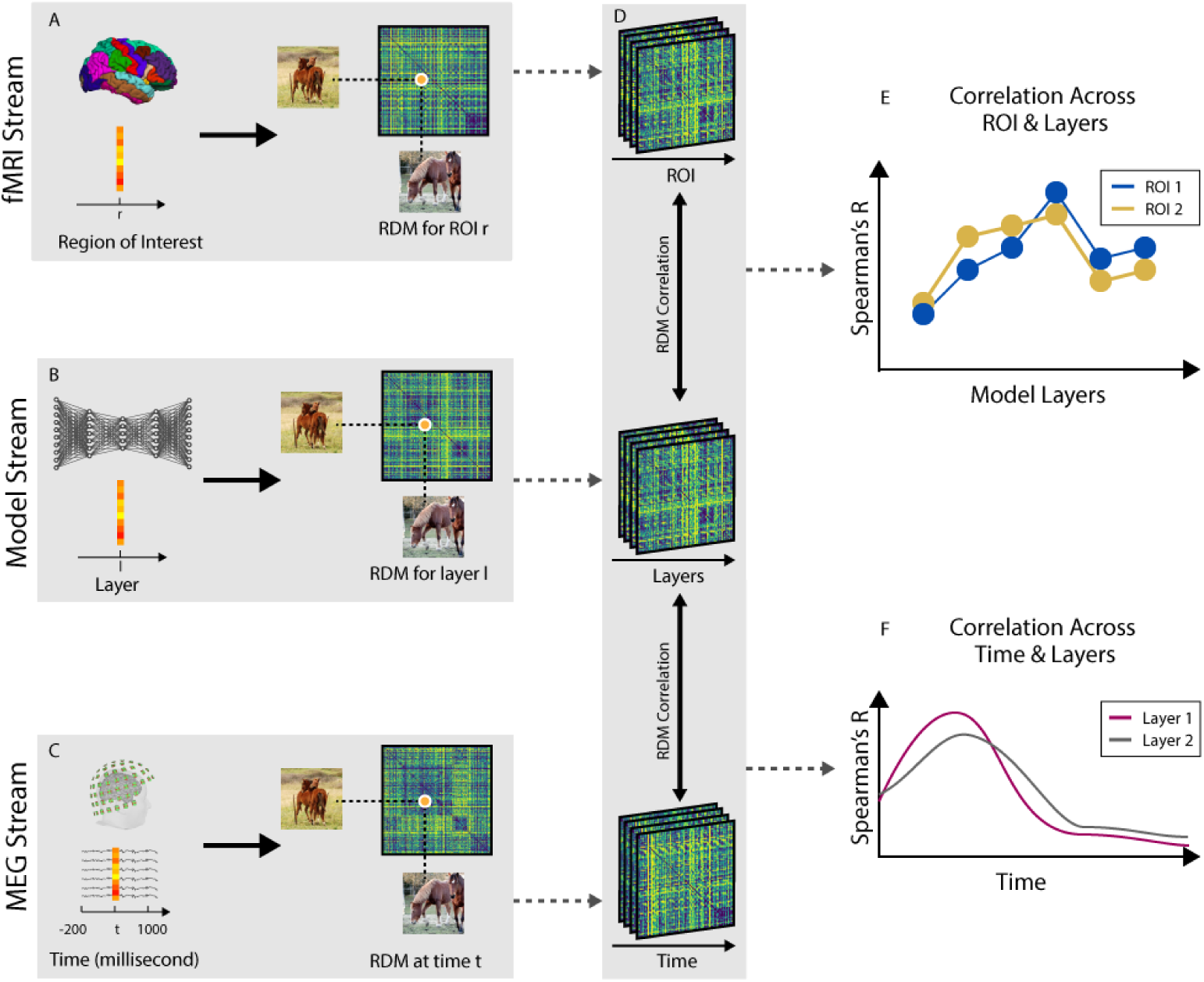
Representational similarity analysis to compare fMRI, MEG and model representations. (A) fMRI response patterns were extracted from each ROI and pairwise condition-specific dissimilarities (1-Pearson’s R) were computed to create one fMRI RDM per ROI and participant (see Materials and methods section for detail). (B) RDMs for the generative model were computed at each convolutional/deconvolutional block after feeding 156 images to the computational model. (C) MEG data consists of time-series data with 306 channels and 1200 time points (milliseconds) per trial. For each condition, we extracted a vector of size 306 at each time point as the activity pattern to compute the RDMs using SVM classifiers decoding accuracies (see Materials and methods section for detail). (D) Using RDMs from MEG and fMRI ROIs, we compared (Spearman’s R) them with the RDMs from the computational model to investigate the spatio-temporal correspondences between the human brain and the computational model. (E) Correlations between time-resolved MEG RDMs and computational model RDMs result in a subject-specific signal for each layer across time, which we then average them over subjects.

**Fig 4.**
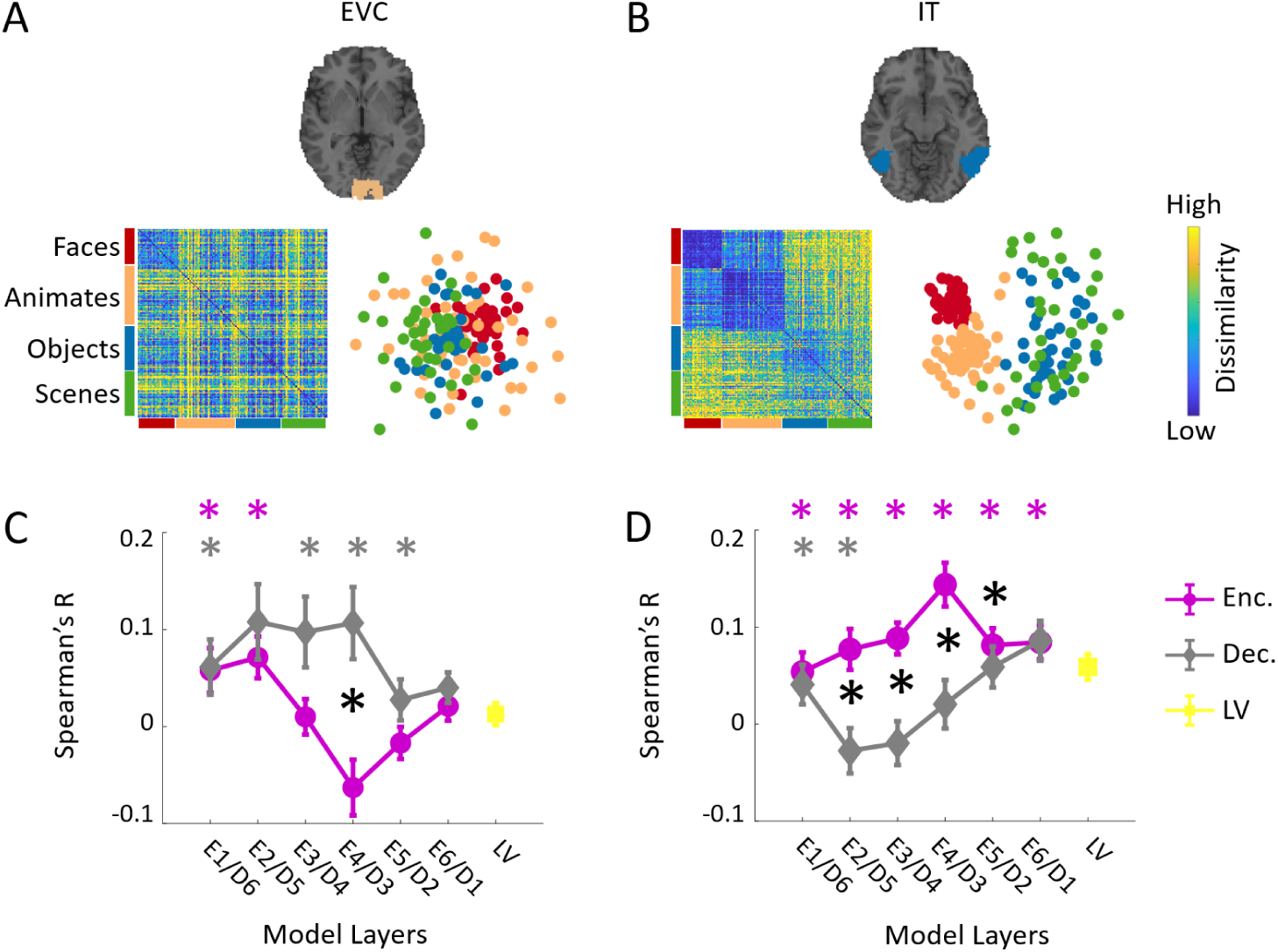
Spatial representational comparisons. (A) Neural representations in early visual cortex (EVC). The subject-averaged EVC RDM matrix, and its 2D multidimensional scaling visualization. (B) Neural representations in inferior temporal area (IT). The subject-averaged IT RDM matrix, and its 2D multidimensional scaling visualization. (C) Encoder, decoder and LV layer RDMs are correlated (Spearman’s R) with subject-specific EVC RDMs. The averaged correlations over subjects with standard error of the mean are depicted. (D) Encoder, decoder and LV layer RDMs are correlated (Spearman’s R) with subject-specific IT RDMs. The averaged correlations over subjects with standard error of the mean are depicted. The color coded (*) above each panel in C-D indicates that the correlation of the corresponding layer is significantly above zero. The black (*) indicates the correlations of the corresponding encoder and decoder layers are significantly different (N=15; two-sided ttests; false discovery rate corrected at *P* < 0.05).

The EVC representational correlations across layers of the encoder (and inverse order of the decoders) demonstrate a decreasing trend, while IT representational correlations across model layers progressively increase. Notably, early processing layers of the model (E1/D6) and late processing layers (E6/D1) show similar correlations both for EVC and IT. However, middle layers correlations are significantly different when comparing encoder and decoder in both ROIs (N=15; two-sided ttests; false discovery rate corrected at *P* < 0.05). Further, the correlations of EVC is stronger with the decoder than encoder whereas the correlations of IT is higher for the encoder than the decoder. This indicates the reconstructing processes are more similar to detail representations in EVC than IT.

### The generative model unfolds the temporal dynamics of brain feedforward and feedback representations

The visual information traverses a hierarchy of regions in the visual cortex and evolves over time rapidly. While the structure of our proposed model does not directly delineate the human brain temporally, it has a clear sequential structure which temporally unfolds the feedforward and feedback sequences in the visual cortex. That is, information flows layer to layer and evolves from image low level features to higher level latent concepts (feedforward sweep) and then from this high level latent code sequentially the image level information is reconstructed (feedback sweep). To test the hypothesis that the proposed model indeed temporally mirrors ventral visual stream dynamics, we compared the representations of time-resolved MEG data acquired in a visual recognition experiment with the layer representations in the encoder and decoder sub-networks. From each participant’s data, we first extracted the MEG sensor measurements for each image condition from −200ms to 1000ms (with 1 ms resolution) relative to image onset. Then we computed dissimilarities (SVM classifiers decoding performances, see Materials and methods section for details) between evoked MEG pattern vectors of each pair of images and created time resolved representational dissimilarity matrices (RDMs) for each individual. We also fed the images to the generator and extracted layer-specific activations for each image condition. Then by computing the pairwise distances (1-Pearson’s R) of image evoked layer activation patterns, we created layer-specific RDMs (see Figure 3 and Materials and methods section for details). Then we correlated layer-specific model RDMs with time-resolved subject specific MEG RDMs (Spearman’s R) resulting in correlation time series for each layer of the model. Figure 5A-B show the subject averaged correlation time series for layers of encoder and decoder sub-networks, respectively. Our results showed that all layers of the proposed model were representationally similar to human brain activity patterns, indicating that the model captures evolution of brain visual representations over time (N=15;permutation tests; cluster definition threshold *P* < 0.05; cluster significance threshold *P* < 0.01). Next, we investigated whether the hierarchy of the layered architecture unfolds the temporal dynamics of encoding (feedforward) and decoding (feedback) visual processes in the brain. Specifically, we examined the relationship between hierarchy of model layers and the peak latency of the correlation time series (Figure 5C). Consistent with previous works [29], the first peak latency of correlation time courses relating MEG and the encoder representations increased with the hierarchy of the encoder layers (Spearman’s *R* = 0.78, *P* << 0.0001). The inspection of peak latencies in the decoder time series depicted in Figure 5B revealed a progressively increasing pattern from D1 to D6 (Spearman’s *R* = 0.84, *P* << 0.0001). That is, the peak latency in D1 occurs at 151ms and over the decoder layers progressively the peak latency increases, the peak latency of D6 occurs at 204ms. Given that the decoder later layers are reconstructing the visual stimuli and thus closer to image level feature space, these temporal dynamics may explain the temporal dynamics of feedback information sent down the ventral stream. We further identified a salient second peak in layers E1 to E5 which their peak latencies negatively correlated with layers order in the Encoder (Spearman’s *R* = −0.57, *P* << 0.0001) and they are significantly later than the corresponding layers in the decoder (all peak latency analyses are based on permutation based bootstrapping; N=15; two-sided hypothesis tests; *P* << 0.0001; Bonnferoni corrected). Again confirming a later dynamics of representational similarities between these layers and the brain, possibly indicating a second sweep of visual information processing.

**Fig 5.**
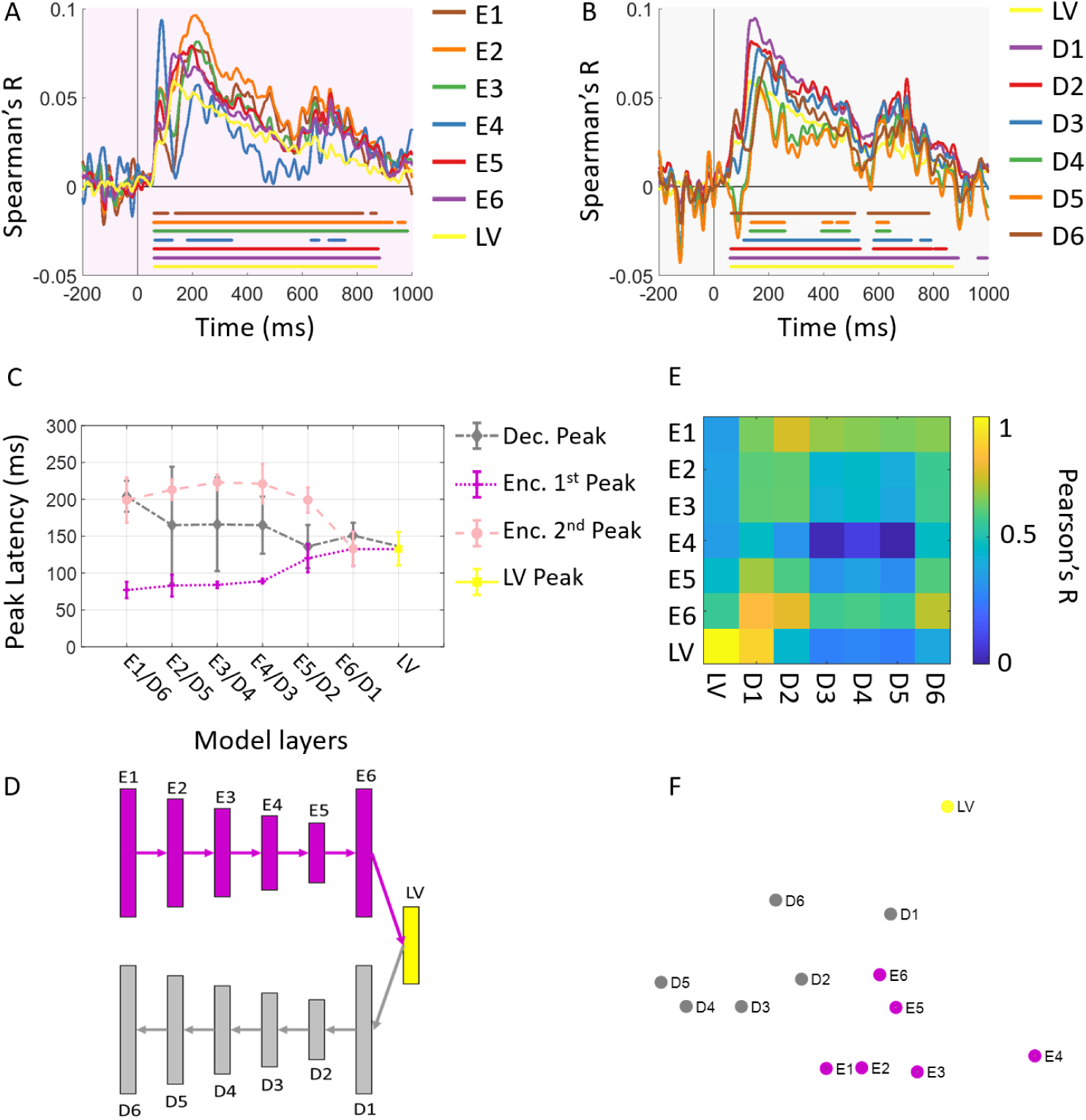
Temporal representational comparisons. (A) Encoder and MEG representational comparison. We correlated the encoder layer RDMs with subject-specific time-resolved MEG RDMs resulting in fifteen correlation time courses. We then averaged these time courses over participants. (B) Decoder and MEG representational comparison. Correlation of the decoder layer RDMs and time-resolved MEG RDMs. The color-coded lines below the curves show the time points when the correlations are significantly above zero (N=15; permutation tests; cluster definition threshold *P* < 0.01; cluster threshold *P* < 0.05). (C) Peak latency for encoder and decoder. The encoder have significantly earlier peak latency across all layers (*P* = 0.014). Error bars are expressed in standard error of the mean. (D) The architecture of the models with layers’s label corresponding to (C). (E) The visualization of relationships between model layers representations.The matrix of RDM correlations between encoder and decoder layers is depicted. Each matrix entry compares two RDMs indexed by corresponding row and column in terms of Pearson’s R. (F) The multidimensional scaling visualization of the RDMs relationships.

In the next step, we investigated the representational relationships among the encoder and the decoder layers of the model. To this end, we first computed the pairwise correlations between all encoder and decoder layer RDMs. The matrix depicted in Figure 5E summarizes these pairwise correlations and reveals which representations across model layers are similar or dissimilar. Figure 5F visualizes these relationships with multi dimensional scaling (MDS) method. Visual inspection of this matrix manifests firstly the dissimilarity of latent layer representation from encoder and decoder layers, secondly similarity of late layer of encoder (E6) and early layer of the decoder (D1); and also similarity of early layer of decoder (E1) and late layer of the decoder (D6), thirdly the dissimilarity of middle layers of encoder and decoder indicating the difference in the representations of encoding (feedforward) and decoding (feedback) processes.

To obtain a more clear picture of brain-model temporal dynamics relationships across encoder and decoder sub-networks, we compared the correlation time courses corresponding to the encoder and decoder layers with the same level of processing in Figure 6A. The correlation time course of latent layer is depicted separately. Figure 6B depicts the corresponding model layers RDMs and their 2-dimensional MDS visualizations. As demonstrated in Figure 5, firstly the low level feature processing layer of encoder (E1) and decoder (D6) follow a notably similar dynamics. Further, the high level feature processing layers of encoder (E6 and E5) also depict a similar temporal dynamics with high level feature processing layers of the decoder (D1 and D2, respectively).

**Fig 6.**
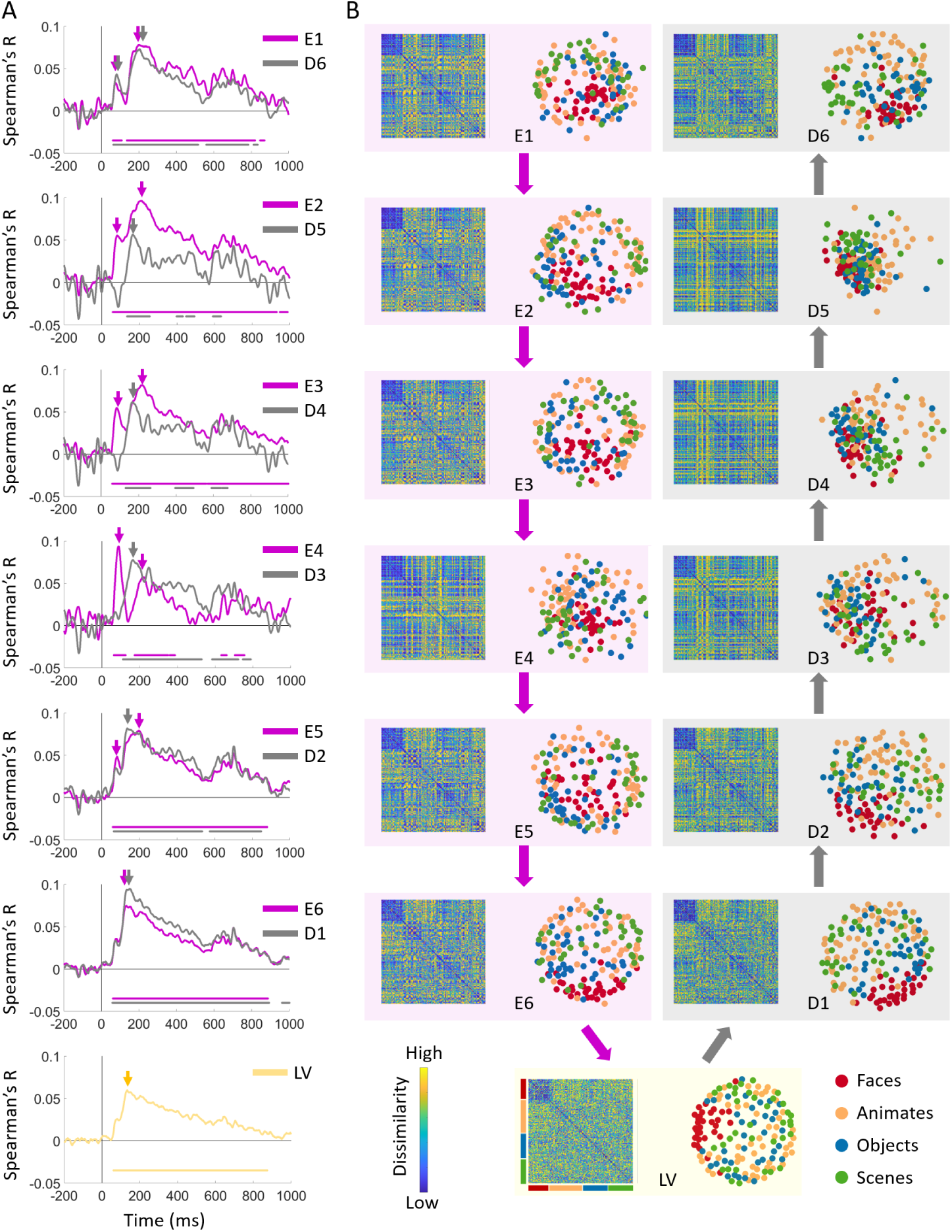
Comparisons of encoder and decoder representational dynamics. (A) Comparison of correlation time series of the encoder and decoder layers with the same level of processing. The color-coded lines below the curves show the timepoints when the correlations are significantly above zero (N=15; permutation tests; cluster definition threshold *P* < 0.01; cluster threshold *P* < 0.05). (B) The model RDMs and their corresponding MDS visualizations.

However, the dynamics are explicitly different when middle level layers are compared (i.e. E2 vs.D5, E3 vs. D4, E4 vs. D3). This is also evident from comparison of the MDS visualizations of RDMs in each row. The peak latencies of the correlation time courses are marked with color-coded arrows. Consistent with Figure 5C, the peak latencies of low and high level processing layers coincide around the same time, whereas the peaks of mid-level feature processing layers of the decoder occurs between the first and second peaks of the corresponding encoder time courses.

Together, comparison of MEG temporal representations with the encoder and decoder sub-networks of our proposed model segregated the brain representational dynamics that transforms the low level visual features to high level categorical semantics and the inverse functional processes that reconstructs low level features from the high level code. These two identified dynamics of processing can be associated with feedforward and feedback sweeps along ventral visual stream.

## Discussion

Using a generative model, we dissected the dynamics of processes in the ventral visual pathway into two temporally distinct stages: the initial sweep depicts neural representational similarity with the hierarchy of representations along the encoder sub-network which transforms visual features into a latent representation (feedforward sweep); and the subsequent sweep shows neural representational similarity with the decoder sub-network which layer by layer reconstructs visual details from the latent representational code (feedback sweep).

As demonstrated in Figure 5A-C, the temporal representational similarities of the encoder sub-network replicates previous findings showing hierarchical representational similarities between the visual brain and the feedforward DNNs that are trained on object recognition tasks [21, 26, 27, 29, 32, 39]. Specifically, it has been demonstrated that object recognition DNNs develop internal representations that are hierarchically similar to brain representations in early visual cortex [39], area V4 (a mid-level region), and IT cortex (a high level categorical region) along ventral visual pathway in primates [21, 26] and humans [27, 29, 32]. Unlike the object recognition DNNs, the encoder sub-network in our model is trained in an unsupervised manner, i.e., the information regarding image classes (e.g. a car, a basketball) were not provided to this network during training. This is a critical finding, as it shows supervision is not required for the development of this hierarchical architecture observed in DNNs and the brain. Further, comparing the representational dynamics of the decoder sub-network and the visual brain revealed a temporally subsequent hierarchy of processing (Figure 5B-C) which progressively builds visual detail from the latent representation. This indicates that beyond feedforward sweep, brain visual processes demonstrate similarities with the reconstruction function implemented in the decoder sub-network. This finding contributes to unraveling the algorithmic functional role of feedback processes in the visual cortex.

Finally, representational comparison of two brain regions at the beginning and end of ventral visual pathway, EVC and IT, with the encoder and decoder subnetworks (Figure 6) revealed the encoding processes in the feedforward sweep develops a categorical representations similar to IT and the reconstructing processes in feedback sweep evolves into detail representations similar to EVC.

## Materials and methods

Our study consists of two components: (i) The computational model and (ii) the MEG/fMRI data from human participants, both of which are analyzed and compared in the result section. We focus on the ventral visual pathway, hence, we acquire both human brain and computational model neuronal activations on visual centric tasks. In this section, we will describe the computational model, SP (Super Category) image dataset, and MEG/fMRI data acquisition and analysis.

### Neuroimaging experiments

The fMRI and MEG data used in this study has been published in [13] previously and is publicly available at http://twinsetfusion.csail.mit.edu/. In this section, we briefly describe the experiment design, data acquisition and analysis.

### Participants

Brain data were acquired from fifteen right-handed healthy participants with normal or corrected to normal vision in two separate experiments (MEG and fMRI). Participants (9 females, 27.87 ± 5.17 years old) signed an informed consent form and were compensated for their participation. Both experiments were conducted in accordance with the Declaration of Helsinki and approved by the Institutional Review Board of Massachusetts Institute of Technology.

### Stimulus set and experimental design

The stimulus set consists of 156 natural images of four distinct visual categories: (i) Faces, (ii) Animates (animals and people), (iii) Objects, (iv) Scenes [13]. The participants viewed images presented for 0.5 sec (with 2.5 sec inter stimulus interval (ISI) in fMRI sessions and 0.7 to 0.8 sec ISI in MEG session) at the center of the screen at 6° visual angle. Functional MRI data were acquired in two sessions (11-15 runs in total) and MEG data were acquired in one session of 25 runs. Images were presented once in each run and in random order. The participants were performing a vigilance task of oddball detection.

### fMRI data acquisition and analysis

The fMRI experiment was conducted at the Athinoula A. Martinos Imaging Center at MIT, using a 3 T Siemens Trio scanner with 32-channel phased-array head coil. Each imaging session started with acquiring structural images using a standard T1-weighted sequence (176 sagittal slices, FOV = 256 mm2, TR = 2530 ms, TE = 2.34 ms, flip angle = 9°) and then 5–8 runs of 305 volumes of functional data (11–15 runs across the two sessions). Gradient-echo EPI sequence was used for functional data acquisition (TR = 2000 ms, TE = 29 ms, flip angle = 90°, FOV read = 200 mm, FOV phase = 100%, bandwidth 2368 Hz/Px, gap = 20%, resolution = 3.1 mm isotropic, slices = 33, ascending interleaved acquisition). For preprocessing of fMRI data, we used SPM software. The preprocessing of functional data included slice-time correction, realignment and co-registration to the first session T1 structural scan, and normalization to the standard MNI space. For multivariate analysis, we did not smooth the data.

We used general linear modeling (GLM) to estimate fMRI responses to the 156 stimuli. The events including stimuli conditions and nulls were modeled with event onsets and impulse response function. Further, the motion and run regressors were included in the GLM. Then we convolved the defined regressors with the hemodynamic response function and estimated the beta-values for each stimulus condition. Then by contrasting each image condition with the explicitly defined null condition, we obtained t-mpas per image condition for each participant. For the current study, we investigated two anatomically defined [36] regions of interest (ROIs) along the ventral visual stream, early visual cortex (EVC) and inferior temporal cortex (IT).

We used multivariate analysis and computed pairwise dissimilarities between 156 image specific fMRI responses using 1-Pearson correlation distances and constructed a 156 × 156 representational dissimilarity matrix (RDM) per participant per ROI (EVC and IT). In detail, we extracted t-value patterns corresponding to each image condition from each region of interest, arranged them into vectors. Then we calculated the pairwise distances of the 156 image specific vector patterns. With this process, we obtained a 156 x156 RDM per ROI for each participant.

### MEG data acquisition and analysis

The MEG experiment was conducted at the Athinoula A. Martinos Imaging Center at MIT, using a 306-channel Elekta neuromag TRIUX system with sampling rate of 1 kHz. The acquired data were filtered by a 0.03 to 330 Hz band-pass filter. We measured the participants’ head position prior and during the recording with 5 coils attached to their head. We then applied a maxfilter for temporal source space separation and head movements correction [40, 41]. For preprocessing of MEG data, we used Brainstorm software [40]. We extracted trials from −200 ms to 1000 ms with respect to image onset. We then removed the baseline mean and smoothed the data with a 30 HZ low-pass filter. For each participant, we obtained 25 trials per image condition. We employed multivariate pattern analysis to compute the dissimilarity relations between image conditions [10–13, 29, 42, 43]. At each time point t, we arranged MEG sensor measurements of each image condition into pattern vectors of 306 x N dimension, where N denotes the number of trials per condition. We then randomly assigned the trials of each condition into 8 bins and subaveraged the trials within each bin to overcome computational complexity and reduce noise. Support vector machine classifiers were trained on the subaveraged MEG pattern vectors of each pair of images at each time point to discriminate the pairs. The performance of the classifier in discriminating each pair of images with leave-one-out cross validation procedure was used as the dissimilarity measure between the pairs to populate a 156 × 156 representational dissimilarity matrix (RDM) at each time point.The rows and columns of the RDM are indexed by the image conditions and each matrix element indicates the dissimilarity of the corresponding image conditions based on MEG measurements of the specific time point.

### Computational Model Architecture and Training

Previous work revealed that deep convolutional neural networks trained on object recognition develop similar representations akin to the hierarchical processes along ventral visual stream [3, 26–32]. However, there are abundant number of feedback connections in ventral visual stream and therefore, these feedforward neural network models may not fully represent the complex visual processes in the ventral visual pathway.

Here, we aim to investigate whether a deep generative model trained to map images to a latent code and then reconstruct the images from the features encoded in the latent space can reveal similar representations as feedforward and feedback processes in the ventral visual pathway. With this aim, we developed a deep generative autoencoder neural network model using adverserial autoencoder (AE) framework [34, 35].

AEs consist of two sub-networks: (i) The encoder takes a given data and outputs a lower representation of the data (latent code) and (ii) the decoder takes the latent code and aims to reproduce the input data. Alternatively, the GAN framework is a min-max adversarial game between two distinct neural networks: (i) The generator(*G*), aims at generating synthetic data by learning the distribution of the real data and (ii) the discriminator (*D*), aims at distinguishing the generator’s fake data from real data. The generator uses a function *G*(*z*) that maps samples *z* from the prior *p*(*z*) (normal distribution) to the data space *p*(*x*). *G*(*z*) is trained to maximally confuse the discriminator into believing that samples it generates come from the data distribution. The solution to this game can be expressed as following [35]:

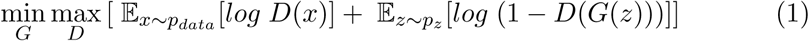

We chose an AE architecture because we hypothesize that the encoder embodies similar neuronal characteristics as the image classification DNNs and thereby could resemble the human brain feedforward representations.

Alternatively, we hypothesize that the decoder part of the AE architecture which generates the image from the latent space code would encompass the neuronal representations similar to feedback processes in the human visual brain.

Our generative autoencoder model architecture consists of a total of 13 layers: (i) the encoder consists of 6 layers; 5 convolutional layers and one fully-connected layer, (ii) the decoder consists of 6 layers; 5 deconvolutional layers and one fully-connected layer, and (iii) lastly one fully-connected layer representing the latent code layer The latent code vector z captures high level representation of the data distribution. The discriminator architecture consists of a total of three fully connected layers (Figure 1B).

### Super Category (SC) Dataset

In computer vision literature, deep neural network models are usually trained to optimize category specific recognition performance on large scale datasets.

However, the human brain learns to develop high-level representations for categories across multiple recognition tasks (eg faces, objects, scenes, etc). Indeed, years of cognitive neuroscience has demonstrated brain regions which are functionally respond preferentially to one of these categories compared to others (eg. Fusiform gyrus, IT area, Parahipocampal cortex, …). Therefore, to train our proposed model we put together a super category (SC) data set consisting of 1,980,00 images from four equally distributed distinct categories: (i) Animals, (ii) Objects, (iii) Scenes, and (iv) Faces. Images from the Faces category were acquired from the VGGFaces2 dataset [44], Objects and Animals categories were from the ImageNet dataset [45], and Scenes categories were from the Places356 dataset [46]. To make all classes equal, we have randomly sampled 495,000 images per class. During computational model training and testing, each image was preprocessed through a pipeline: (i) Images were resized to 224 × 224, and (ii) normalized from 0 to 225 to −1 to 1 range values.

Lastly, the neuronal representations for the generative AE model were computed at each convolutional/deconvolutional block after feeding 156 images used in the neuroimaging experiments to the encoder of the computational model. Please note this image set was not used in training the model. We employed the vectorized model activity patterns of each convolutional/deconvolutional block to compute dissimilarity distances (1-Pearson’s R) for each pair of imagesand create an RDM per model layer (Figure 3B).

### Representational similarity analysis to relate the brain and model representations

We used representational similarity analysis (RSA) [37, 38] to map MEG measurements, fMRI responses, and model activation patterns into a common space where they are directly comparable.

RSA transforms the stimulus-specific response patterns into a representational space by creating matrices summarizing pairwise distance relationships of the response patterns (i.e. defined as the correlational distance, or a classifier performance in discriminating two conditions). The matrix capturing these pairwise dissimilarity measures is called representational dissimilarity matrix (RDM).

To relate the spatio-temporal dynamics of neural representations in the human brain with our proposed model, we computed the similarity (in terms of Spearman’s R) of fMRI ROI RDMs and time-resolved MEG RDMs with our computational model RDMs (Figure 3D).

Correlations between subject-specific time-resolved MEG RDMs and computational model layer RDMs result in a signal for each layer per participant across time (Figure 3F). While, correlations between subject-specific fMRI ROI RDMs and computational model layer RDMs result in subject-specific correlation values per layer (Figure 3E). To account for different levels of noise in brain ROIs, we estimated the noise ceiling in EVC and IT [47, 48] and normalized the fMRI ROI and model correlations with the corresponding noise ceiling [49]. Then the correlation time series (for MEG/Model comparisons) or correlation values (for fMRI/Model comparisons) were averaged over participant and tested against zero for statistical significance.

## Statistical tests

We used nonparametric statistical test methods which make no assumptions on the distribution of the data [50, 51]. For statistical inference on the correlation time series, we used permutation-based cluster-size inference with null hypothesis of zero. For statistical assessments of peak latencies, we bootstrapped the subject-specific correlation time series for 1000 times to estimate an empirical distribution over peak latencies [12, 29, 43].

## Acknowledgments

We would like to thank Western BrainsCAN for the generous support of this research. Computational modeling was conducted on Compute Canada resources.

## Notes

### Competing Interest Statement

The authors have declared no competing interest.

## References

1. Epstein R, Kanwisher N. A cortical representation the local visual environment. Nature. 1998;392(6676):598–601. doi: 10.1038/33402.

2. Epstein R, Harris A, Stanley D, Kanwisher N. The parahippocampal place area: Recognition, navigation, or encoding? Neuron. 1999;23(1):115–125. doi: 10.1016/S0896-6273(00)80758-8.

3. Martin Cichy R, Khosla A, Pantazis D, Oliva A. Dynamics of scene representations in the human brain revealed by magnetoencephalography and deep neural networks. NeuroImage. 2017;153:346–358. doi: 10.1016/j.neuroimage.2016.03.063.

4. Lowe MX, Rajsic J, Ferber S, Walther DB. Discriminating scene categories from brain activity within 100 milliseconds. Cortex; a journal devoted to the study of the nervous system and behavior. 2018;106:275–287. doi: 10.1016/j.cortex.2018.06.006.

5. Henriksson L, Mur M, Kriegeskorte N. Rapid Invariant Encoding of Scene Layout in Human OPA. Neuron. 2019;103(1):161 – 171.e3. doi: https://doi.org/10.1016/j.neuron.2019.04.014.

6. Kanwisher N, McDermott J, Chun MM. The fusiform face area: A module in human extrastriate cortex specialized for face perception. Journal of Neuroscience. 1997;17(11):4302–4311. doi: 10.1523/JNEUROSCI.17-11-04302.1997.

7. Dobs K, Isik L, Pantazis D, Kanwisher N. How face perception unfolds over time. Nature Communications. 2019;10(1):1258. doi: 10.1038/s41467-019-09239-1.

8. Grill-Spector K, Kushnir T, Hendler T, Edelman S, Itzchak Y, Malach R. A sequence of object-processing stages revealed by fMRI in the human occipital lobe. Human brain mapping. 1998;6(4):316–328.

9. Grill-Spector K, Kourtzi Z, Kanwisher N. The lateral occipital complex and its role in object recognition. Vision Research. 2001;41(10):1409–1422. doi: https://doi.org/10.1016/S0042-6989(01)00073-6.

10. Cichy RM, Pantazis D, Oliva A. Resolving human object recognition in space and time. Nature Neuroscience. 2014;17(3):455–462. doi: 10.1038/nn.3635.

11. Isik L, Meyers EM, Leibo JZ, Poggio T. The dynamics of invariant object recognition in the human visual system. Journal of Neurophysiology. 2014;111(1):91–102. doi: 10.1152/jn.00394.2013.

12. Mohsenzadeh Y, Qin S, Cichy RM, Pantazis D. Ultra-rapid serial visual presentation reveals dynamics of feedforward and feedback processes in the ventral visual pathway. eLife. 2018;7. doi: 10.7554/eLife.36329.

13. Mohsenzadeh Y, Mullin C, Lahner B, Cichy RM, Oliva A. Reliability and Generalizability of Similarity-Based Fusion of MEG and fMRI Data in Human Ventral and Dorsal Visual Streams. 2019;doi: 10.3390/vision3010008.

14. Goodale MA, Milner AD. Separate visual pathways for perception and action. Trends in neurosciences. 1992;15(1):20–25. doi: 10.1016/0166-2236(92)90344-8.

15. Ungerleider LG, Haxby JV. ‘What’ and ‘where’ in the human brain. Current opinion in neurobiology. 1994;4(2):157–165. doi: 10.1016/0959-4388(94)90066-3.

16. Li P, Zhu S, Chen M, Han C, Xu H, Hu J, et al. A motion direction preference map in monkey V4. Neuron. 2013;78(2):376–388. doi: 10.1016/j.neuron.2013.02.024.

17. DiCarlo JJ, Maunsell JHR. Anterior inferotemporal neurons of monkeys engaged in object recognition can be highly sensitive to object retinal position. Journal of neurophysiology. 2003;89(6):3264–3278. doi: 10.1152/jn.00358.2002.

18. Chakravarthi R, Carlson TA, Chaffin J, Turret J, VanRullen R. The temporal evolution of coarse location coding of objects: Evidence for feedback.; 2014.

19. Serre T, Oliva A, Poggio T. A feedforward architecture accounts for rapid categorization. Proceedings of the National Academy of Sciences. 2007;104(15):6424–6429. doi: 10.1073/pnas.0700622104.

20. DiCarlo JJ, Zoccolan D, Rust NC. How does the brain solve visual object recognition; 2012.

21. Yamins DLK, Hong H, Cadieu CF, Solomon EA, Seibert D, DiCarlo JJ. Performance-optimized hierarchical models predict neural responses in higher visual cortex. Proceedings of the National Academy of Sciences. 2014;111(23):8619–8624. doi: 10.1073/pnas.1403112111.

22. Tang H, Kreiman G. In: Recognition of occluded objects. Singapore: Springer-Verlag; 2017.

23. Rajaei K, Mohsenzadeh Y, Ebrahimpour R, Khaligh-Razavi SM. Beyond core object recognition: Recurrent processes account for object recognition under occlusion. PLOS Computational Biology. 2019;15(5):e1007001. doi: 10.1371/journal.pcbi.1007001.

24. Kar K, Kubilius J, Schmidt K, Issa EB, DiCarlo JJ. Evidence that recurrent circuits are critical to the ventral stream’s execution of core object recognition behavior. Nature Neuroscience. 2019;22(6):974–983. doi: 10.1038/s41593-019-0392-5.

25. Kietzmann TC, Spoerer CJ, Sörensen LKA, Cichy RM, Hauk O, Kriegeskorte N. Recurrence is required to capture the representational dynamics of the human visual system. Proceedings of the National Academy of Sciences of the United States of America. 2019;116(43):21854–21863. doi: 10.1073/pnas.1905544116.

26. Yamins D, Hong H, Cadieu C, Dicarlo JJ. Hierarchical Modular Optimization of Convolutional Networks Achieves Representations Similar to Macaque IT and Human Ventral Stream. NIPS. 2013;.

27. Khaligh-Razavi SM, Kriegeskorte N. Deep Supervised, but Not Unsupervised, Models May Explain IT Cortical Representation. PLoS Computational Biology. 2014;10(11). doi: 10.1371/journal.pcbi.1003915.

28. Cox DD, Dean T. Neural networks and neuroscience-inspired computer vision;2014. Available from: http://www.ncbi.nlm.nih.gov/pubmed/25247371.

29. Cichy RM, Khosla A, Pantazis D, Torralba A, Oliva A. Comparison of deep neural networks to spatio-temporal cortical dynamics of human visual object recognition reveals hierarchical correspondence. Scientific Reports. 2016;6(1):1–13. doi: 10.1038/srep27755.

30. Cichy RM, Kaiser D. Deep Neural Networks as Scientific Models; 2019.

31. Güçlü U, van Gerven MAJ. Deep neural networks reveal a gradient in the complexity of neural representations across the ventral stream. Journal of Neuroscience. 2015;35(27):10005–10014. doi: 10.1523/JNEUROSCI.5023-14.2015.

32. Mohsenzadeh Y, Mullin C, Lahner B, Oliva A. Emergence of Visual Center-Periphery Spatial Organization in Deep Convolutional Neural Networks. Scientific Reports. 2020;10(1):1–8. doi: 10.1038/s41598-020-61409-0.

33. Cichy RM, Roig G, Andonian A, Dwivedi K, Lahner B, Lascelles A, et al. The Algonauts Project: A Platform for Communication between the Sciences of Biological and Artificial Intelligence; 2019.

34. Makhzani A, Shlens J, Jaitly N, Goodfellow I, Frey B. Adversarial Autoencoders. 2015;.

35. Goodfellow IJ, Pouget-Abadie J, Mirza M, Xu B, Warde-Farley D, Ozair S, et al. Generative Adversarial Nets;. Available from: http://www.github.com/goodfeli/adversarial.

36. Tzourio-Mazoyer N, Landeau B, Papathanassiou D, Crivello F, Etard O, Delcroix N, et al. Automated anatomical labeling of activations in SPM using a macroscopic anatomical parcellation of the MNI MRI single-subject brain. NeuroImage. 2002;15(1):273–289. doi: 10.1006/nimg.2001.0978.

37. Kriegeskorte N. Representational similarity analysis – connecting the branches of systems neuroscience. Frontiers in Systems Neuroscience. 2008;2(NOV):4. doi: 10.3389/neuro.06.004.2008.

38. Kriegeskorte N, Kievit RA. Representational geometry: Integrating cognition, computation, and the brain; 2013.

39. Cadena SA, Denfield GH, Walker EY, Gatys LA, Tolias AS, Bethge M, et al. Deep convolutional models improve predictions of macaque V1 responses to natural images. PLOS Computational Biology. 2019;15(4):1–27. doi: 10.1371/journal.pcbi.1006897.

40. Tadel F, Baillet S, Mosher JC, Pantazis D, Leahy RM. Brainstorm: A User-Friendly Application for MEG/EEG Analysis. Computational Intelligence and Neuroscience. 2011;2011:879716. doi: 10.1155/2011/879716.

41. Taulu S, Simola J. Spatiotemporal signal space separation method for rejecting nearby interference in MEG measurements. Physics in Medicine & Biology. 2006;51(7):1759.

42. Carlson T, Tovar DA, Alink A, Kriegeskorte N. Representational dynamics of object vision: The first 1000 ms. Journal of Vision. 2013;13(10):1–1. doi: 10.1167/13.10.1.

43. Pantazis D, Fang M, Qin S, Mohsenzadeh Y, Li Q, Cichy RM. Decoding the orientation of contrast edges from MEG evoked and induced responses. NeuroImage. 2018;180:267–279. doi: 10.1016/j.neuroimage.2017.07.022.

44. Cao Q, Shen L, Xie W, Parkhi OM, Zisserman A. VGGFace2: A dataset for recognising faces across pose and age;. Available from: http://www.robots.ox.ac.uk/.

45. Russakovsky O, Deng J, Su H, Krause J, Satheesh S, Ma S, et al. ImageNet Large Scale Visual Recognition Challenge. International Journal of Computer Vision. 2015;115(3):211–252. doi: 10.1007/s11263-015-0816-y.

46. Zhou B, Lapedriza A, Khosla A, Oliva A, Torralba A. IEEE Transactions on Pattern Analysis and Machine Intelligence Places: A 10 million Image Database for Scene Recognition. 1109;doi: 10.1109/TPAMI.2017.2723009.

47. Nili H, Wingfield C, Walther A, Su L, Marslen-Wilson W, Kriegeskorte N. A Toolbox for Representational Similarity Analysis. PLOS Computational Biology. 2014;10(4):1–11. doi: 10.1371/journal.pcbi.1003553.

48. Oliva A. In: Gist of the Scene. vol. 696; 2005. p. 251–256.

49. Khaligh-Razavi SM, Cichy RM, Pantazis D, Oliva A. Tracking the Spatiotemporal Neural Dynamics of Real-world Object Size and Animacy in the Human Brain. Journal of Cognitive Neuroscience. 2018;30(11):1559–1576. doi: 10.1162/jocna01290.

50. Maris E, Oostenveld R. Nonparametric statistical testing of EEG- and MEG-data. Journal of Neuroscience Methods. 2007;164(1):177–190. doi: https://doi.org/10.1016/j.jneumeth.2007.03.024.

51. Pantazis D, Nichols TE, Baillet S, Leahy RM. A comparison of random field theory and permutation methods for the statistical analysis of MEG data. NeuroImage. 2005;25(2):383–394. doi: 10.1016/j.neuroimage.2004.09.040.

